# Comparing the Impact of Systemic Pituitary Adenylate-Cyclase-Activating Polypeptide (PACAP) and Calcitonin Gene-Related Peptide (CGRP) on Motion-induced Nausea and Balance Behaviors in Mice

**DOI:** 10.1101/2024.10.17.618743

**Authors:** Shafaqat M. Rahman, Abigail Dweh, Anne E. Luebke

## Abstract

Pituitary adenylate-cyclase-activating polypeptide (PACAP), particularly its dominant isoform PACAP-38, is implicated in migraine and represents a promising therapeutic target. We investigated if intraperitoneally delivered (IP) PACAP-38 impacts motion-induced nausea, postural sway, and imbalance in C57BL/6J wildtype mice using the motion-induced thermoregulation, center of pressure (CoP), rotarod, and balance beam assays. We also assessed systemic Calcitonin Gene-Related Peptide’s (CGRP) effects on these behaviors in parallel. Our findings indicate that IP PACAP-38 significantly disrupts motion-induced thermoregulation in mice, with notable blunting of tail vasodilation responses in both sexes. Additionally, PACAP-38 administration increased postural sway in female mice only and caused balance beam imbalances. Contrary to IP CGRP, IP PACAP-38 did not affect rotarod performance when mice were trained on a dowel with 1.5 cm radius. Our findings provide preclinical evidence supporting a potential role of PACAP-38 in vestibular migraine pathophysiology. Future research will explore if PACAP antagonism can protect against PACAP-38’s effects on nausea and balance behaviors, relevant to treatment of vestibular migraine (VM), especially for patients unresponsive to triptans or CGRP-targeting therapies.

## Introduction

Pituitary adenylate-cyclase-activating polypeptide (PACAP) is a neuropeptide that exists in two isoforms, PACAP-27 and PACAP-38, and is implicated in migraine pathophysiology. PACAP-27 and PACAP-38 are strongly associated with their cognate receptor – the PAC1 receptor (PAC1-R) – and both neuropeptides and PAC1-R are found in central and peripheral nervous system structures [1]. Additionally, both PACAP forms can also bind to the vasoactive intestinal peptide/PACAP receptors VPAC1 and VPAC2. In pertinence to migraine, PACAP-38 is in the trigeminal and sphenopalatine ganglia, hypothalamus, cerebellum, and PACAP co-localizes with calcitonin gene-related peptide (CGRP) in the trigeminal system. PACAP-38 is the dominant isoform of PACAP, representing 90% of PACAP circulation [2, 3], and is much further characterized in migraine than PACAP-27.

Recent breakthroughs in migraine treatment have focused on antagonizing calcitonin gene-related peptide (CGRP), and now a strong focus is on evaluating PACAP antagonism [2, 4]. CGRP and PACAP are involved in migraine, as i) infusion of CGRP or PACAP can cause migraine in migraine-sensitive patients [3, 5], ii) CGRP-signaling blockade has resulted in numerous FDA approved therapies [6, 7], and iii) the PACAP-targeting antibody, Lu G0922, is being assessed in clinical trials, with recent results suggesting Lu G0922 effectively inhibits PACAP-38 induce cephalic vasodilation [8], and a high dose of Lu G0922 demonstrates a 2.0 day reduction in monthly migraine days (MMD) in patients with episodic or chronic migraine [9]. Presently, a Phase 2 clinical trial of Lu AG09222 to determine optimal dosing, titled PROCEED (clinical trial identifier: NCT06323928), is currently ongoing.

Preclinical studies on allodynia and light-aversion in mice suggest that CGRP and PACAP act independently, and that antagonizing PACAP may be a therapeutic target for non-responders to CGRP-targeting therapy [10, 11]. In addition, vestibular migraine (VM) is a form of migraine with diagnostic criteria of the consensus document of the International Bárány Society for Neuro-Otology and the International Headache Society, that defines VM to be the combination of typical signs and symptoms of migraine with the vestibular symptoms (5 min to 72 h of duration) of no other potential cause [12, 13]. And even though VM accounts for 7% of patients seen in dizziness clinics and 9% of patients seen in headache clinics it is still underdiagnosed. Currently, research on CGRP/PACAP antagonism is limited in vestibular migraine (VM) therapy. Vestibular migraine can occur with or without headache, and VM patients exhibit an enhanced motion sensitivity that translates to motion sickness, vertigo, dizziness, and postural sway [14–17]. A recent research report indicated that as many as 80% of VM patients experienced relief when treated with CGRP antagonists [18], and another recently published randomized trial for galcanezumab for vestibular migraine showed this anti-CGRP monoclonal was effective in 65-70% of VM patients [19] yet there are patients that are non-responders suggesting other signaling pathways could be targeted in VM.

In this study using the C57BL/6J wildtype mice, we evaluate the effects of the dominant form of PACAP, PACAP-38, on motion-induced nausea, postural sway, and imbalance using the motion-induced thermoregulation, center of pressure, rotarod, and balance beam assays. Both PACAP-38 and CGRP are administered intraperitoneally (IP), with CGRP assessed in parallel to compare with PACAP-38 effects.

## Methods

### Animals

Wildtype C57BL/6J mice were obtained from Jackson Laboratories. Mice were housed under a 12-hour day/night cycle under the care of the University Committee on Animal Resources (UCAR) at the University of Rochester. Mice had *ad libitum* access to necessities (food, water, bedding, and enrichment). A total of 89 mice (47F/42M) were tested in this study, and different groups of mice were allocated for testing motion-induced nausea (n = 16F/16M), postural sway (n = 22F/18M), rotarod (n = 9F/8M), and balance beam (n = 5F/5M). All studies were sufficiently powered. Mice were equilibrated in a testing room with an ambient temperature between 22-23°C for at least 30 minutes prior to testing. Mice were tested between 3 to 7 months of age. Testing occurred from 9:00 am to 5:30 pm.

### Drug administration

All intraperitoneal (IP) injections were performed with a fine 33-gauge insulin syringe. Dulbecco PBS (saline) served as vehicle control and as diluent. IP CGRP was prepared at 0.1 mg/kg (rat ɑ-CGRP, Sigma-Aldrich C0292) and PACAP-38 was prepared at 0.3 mg/kg (Bachem 124123-15-5). Mice were tested on behavioral assays about 20 to 25 minutes after IP injections. Mice were gently handled without the need for anesthesia. All animal procedures were approved by the University of Rochester’s (IACUC) and performed in accordance with the standards set by the NIH.

### Motion-induced thermoregulation

We used a FLIR E60 IR camera (model: E64501) to measure head and tail temperatures of mice before, during, and after a vestibular perturbation (**Fig 1A**). The measurements were conducted over a duration of 45 minutes, following a previously published protocol where we evaluated the effects of IP CGRP with and without migraine blockers on this assay [20]. To summarize, we recorded baseline measurements for five minutes prior to initiating the perturbation (-5 mins ≤ t ≤ 0 mins). The head measurement was between the eye sockets, and the tail temperature measured 1 cm from body or proximal tail. Subsequently, mice were recorded for 20 minutes (0 mins ≤ t ≤ 20 mins) during an orbital rotation at 75 rpm (2 cm orbital displacement). After the perturbation, mice were additionally recorded for 20 minutes to observe the recovery from hypothermia back to baseline (20 min ≤ t ≤ 40 min). FLIR Tools+ was used for analyzing data.

**Figure 1:**
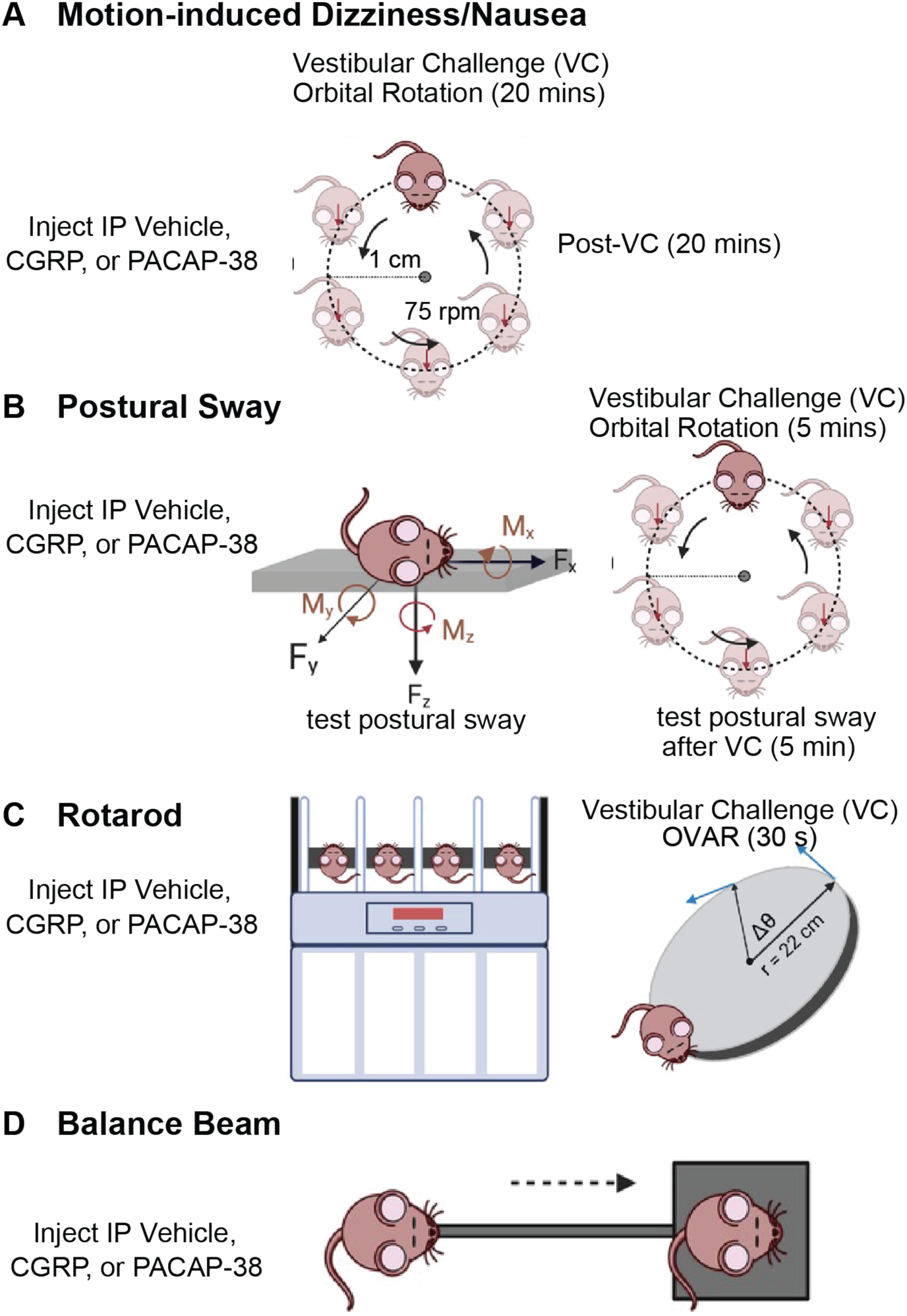
**A.** Mouse surrogate behavior for nausea/motion sickness is to study motion-induced thermoregulation, involving 45-minute head and tail temperature recording consisting of 5-minute baseline, 20 minutes with provocative motion, and a 20-minute recovery period. **B.** Postural sway is evaluated using the center of pressure (CoP) assay, and mice are tested before and after a 5-minute orbital rotation (125 rpm). **C.** Rotarod assesses dynamic balance in which mice travel on an accelerating, rotating dowel, and mice were evaluated before and after off-vertical axis rotation at 60 rpm, 45° tilt for 30 seconds. **D.** Balance beam is another assay for mouse balance and measures the time for mice to traverse from one end of the beam to the other. Mice were tested on behavioral assays after intraperitoneal injections (IP) of vehicle (phosphate buffered-saline), 0.1 mg/kg CGRP, and 0.3 mg/kg PACAP-38 (with exception of rotarod, which did not involve CGRP testing).

We defined Δ tail vasodilatations (°C) as the transient increases in mouse tail temperature in response to perturbation. These were computed by subtracting the baseline tail temperature at time t = 0 minutes from the maximum tail temperature measured during the first 10 minutes of the rotation (0 ≤ t ≤ 10). Additionally, we calculated the magnitude of hypothermia (Δ head temperature) by subtracting the baseline head temperature at time t = 0 minutes from the minimum head temperature recorded throughout the entire experiment (0 mins ≤ t ≤ 40 mins).

### Postural sway testing before and after a 5-minute orbital rotation

Postural sway was evaluated via the center of pressure (CoP) assay (**Fig 1B**) using the AMTI Biomechanics Force platform (model HEX6×6) following the same protocol and setup as previously described in another paper from our group [21]. A quick overview is provided. CoP is represented as a 95% confidence ellipse area that encloses 95% of the CoP trajectory values recorded in a single trial. Each 95% CoP ellipse area can be further decomposed into its major and minor axes, which we also analyzed in this study. About 8 to 10 CoP trials were recorded per mouse in each experiment, and a 10% robust outlier removal (ROUT) was applied to eliminate outlier CoP areas per mouse. After this exclusion, each mouse had a minimum of six CoP areas, which were averaged to compute the mouse’s average 95% confidence ellipse area (cm^2^) in each experimental condition. For each drug treatment (vehicle, CGRP, or PACAP-38), mice were tested for postural sway before and after a 5-minute orbital rotation at 125 rpm (1 cm orbital radius). No individual mice were excluded from the study.

### Rotarod testing before and after off-vertical axis rotation (OVAR)

Dynamic balance was assessed with the Rotarod (Columbus Instruments) setup (**Fig 1C**) with the mouse dowel (r = 1.5 cm), and we used the same protocol as we published in a recent study, with the same two cage rotator that creates the off-vertical axis rotation (OVAR) stimulus [22]. In summary, mice were tasked to maintain balance on the mouse dowel rotating from 5 to 44 rpm at an acceleration step of 2.4 rpm every 4 seconds. Latency to fall (LTF) is measured when mice fall from the dowel. Three days of rotarod testing were performed, with day 1 for training, day 2 for IP vehicle assessment, and day 3 for IP PACAP-38. Twenty minutes after the injection, mice were tested for 3 trials (pre-VC), then stimulated with OVAR (60 rpm, 45° tilt from the vertical) for 30 seconds. Mice were then immediately tested for an additional 3 trials (post-VC).

### Balance beam testing

A specially designed lab balance beam assay was used to measure motor coordination and balance. The balance beam used in these studies was an elevated metal bar that was 80 cm long and 9 mm wide. On one end of the beam, there was an enclosed box with food pellets and a red enrichment house from the animal’s home cage. The animals were placed on the opposite end of the beam, which was the beginning position. In order to entice animals to cross the beam, the enclosed box was darker, and the starting position was well-lit with 60W light (**Fig. 1D**). Mice were taught on the balance beam for three to five trials on the first day. In the days and trials that followed, three trials were conducted with mice positioned on the beam, and the average time required to reach the home box was calculated. Every beam session was captured on an iPhone (model X) as a short movie (∼ 3 min duration). Eash session and trial was subsequently uploaded to BOX for two reviewers who were blinded to examine later. The crossing duration was the amount of time the mouse body took to cross the leading and following yellow markers. All mice received intraperitoneal injections (IP) of either vehicle, CGRP, or PACAP-38, at least 25 minutes before testing. 70% ethanol was used to wipe the beam and the home cage of each animal following testing.

### Data analysis and statistics

All statistical analyses were conducted in GraphPad Prism 10. Three-way repeated measure ANOVAs (RM-ANOVAs) were used to analyze postural sway and rotarod outcomes, two-way mixed-effect models were used to analyze Δ tail vasodilatations and Δ head temperatures, and a two-way RM-ANOVA was used to assess balance beam activity. Group averages of Δ tail vasodilatations, postural sway, and balance beam outcomes are reported as mean ± SEM, and significance was set at p < 0.05 for all analyses.

Analysis of Δ tail vasodilatations involved defining a threshold of 1.5°C to examine the data as a binary outcome. Tail temperature changes equal to or greater than +1.5°C were designated a Δ tail vasodilation and those less than +1.5°C did not meet the criteria and were labeled as diminished tail vasodilatations. Mice were excluded from further testing if their Δ tail vasodilation after vehicle (saline) testing were below 1.5°C. This binary criterion was also used to determine the percentage of mice that exhibited diminished Δ tail vasodilatations after neuropeptide (PACAP-38 or CGRP) treatment. We did not impose a binary outcome to Δ head temperatures.

Second-order curve fitting was used for fitting head temperatures (B2*X^2^ + B1*X + B0) and R^2^ fit were calculated per curve. Head recovery (mins) was approximated by normalizing head temperature at x = 0 mins to y = 0 °C. An x-intercept quadratic model approximated the recovery by detecting x-intercepts passed 20 minutes (constrained at x > 20 minutes, when we stopped the orbital rotation).

For rotarod, each mouse participated in 3 trials, and the max latency to fall (LTF) of the three trials was analyzed between treatment groups.

For balance beam, each mouse underwent three trials, and the results from these trials were averaged to obtain a mean value for each individual mouse. For statistical analysis, the mean values of all mice in the treatment group were then averaged to calculate the overall group mean + SEM. If a mouse fell off the balance beam or stalled a max time of 120 seconds was denoted. We additionally quantified the confidence interval (CI) and coefficient of variation (COV) per treatment group.

## Results

### IP PACAP-38 disrupts motion-induced thermoregulation in C57/B6J mice

Motion-induced thermoregulation was assessed in all mice treated with IP saline as the vehicle (n = 16F/16M), and we divided mice into two groups for testing either IP CGRP (n = 8F/8M) or IP PACAP-38 (n = 8F/8M) testing. **Figure 2A-F** show tail temperature profiles of mice after IP vehicle (**Fig2A-B**) IP CGRP (**2C-D**) or IP PACAP-38 (**2E-F**) for males and females. We computed the delta tail vasodilatations for each mouse in each test and ran a 2-way mixed effects model examining the factors Sex (male vs female) and Treatment (IP Vehicle vs IP CGRP vs IP PACAP-38). Statistical analysis showed effects of Treatment (F (2, 28) = 69.20, p < 0.0001), but no effect of Sex (F (1, 30) = 1.35) or the interactions of these factors (F (2, 28) = 0.42). In agreement with our previous findings [20], Tukey multiple comparison’s test indicates IP CGRP blunted tail vasodilatations compared to their vehicle responses in females (adj. p < 0.0001) and in males (adj. p < 0.0001). Interestingly, IP PACAP-38 also blunted tail vasodilatations in females (adj. p < 0.0001) and in males (adj. p < 0.0001). 100% of females and 75% of males exhibited diminished delta tail vasodilatations (< delta 1.5 °C) with IP CGRP testing, and 100% of females 87.5% of males exhibited diminished delta tail vasodilatations with IP PACAP-38. Biological sex did not exert an effect on tail vasodilatations within the IP vehicle, CGRP, and PACAP-38 treatment groups.

**Figure 2:**
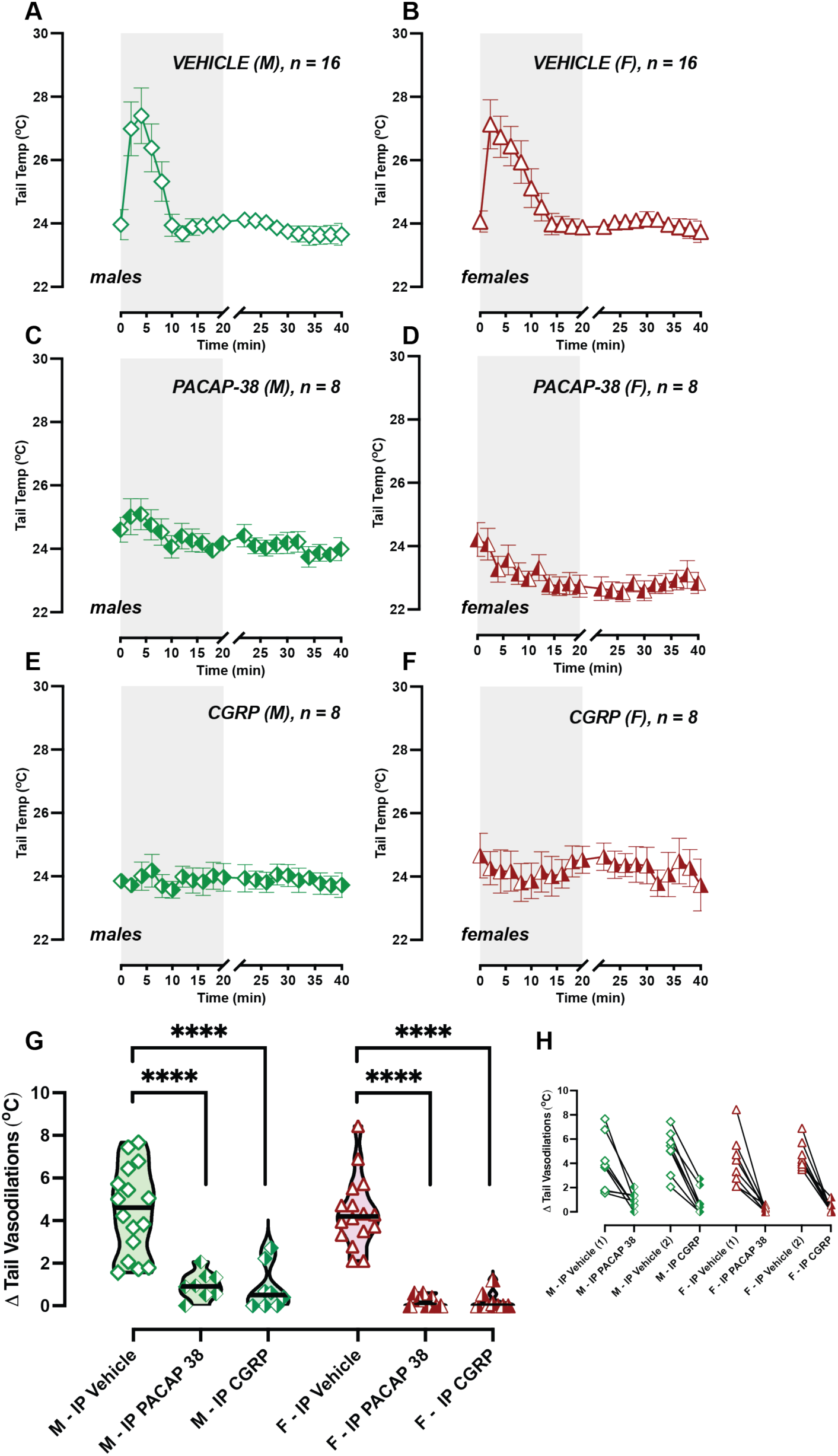
IP PACAP-38 and CGRP disrupt tail vasodilatations to provocative rotation. Tail temperature profiles of C57BL/6J mice during testing are depicted for IP vehicle (**A and B),** IP CGRP (**C and D**), and IP PACAP-38 (**E and F**), with male graphs placed on left and females on right. Symbols are illustrated in the following scheme: green diamonds: male, red triangles: female, open symbol: IP vehicle, right-shaded symbol: CGRP, left-shaded symbol: PACAP-38. Sample sizes are depicted in legend. **G.** Delta tail vasodilatations indicate disrupted thermoregulation after IP CGRP and PACAP-38 (Tukey post hoc, adj. p < 0.0001). **H.** Before/after plot shows intra-animal changes, with Vehicle (1) indicating mice allocated for CGRP tests, and Vehicle (2) indicating PACAP-38 tests.

**Figure 3:**
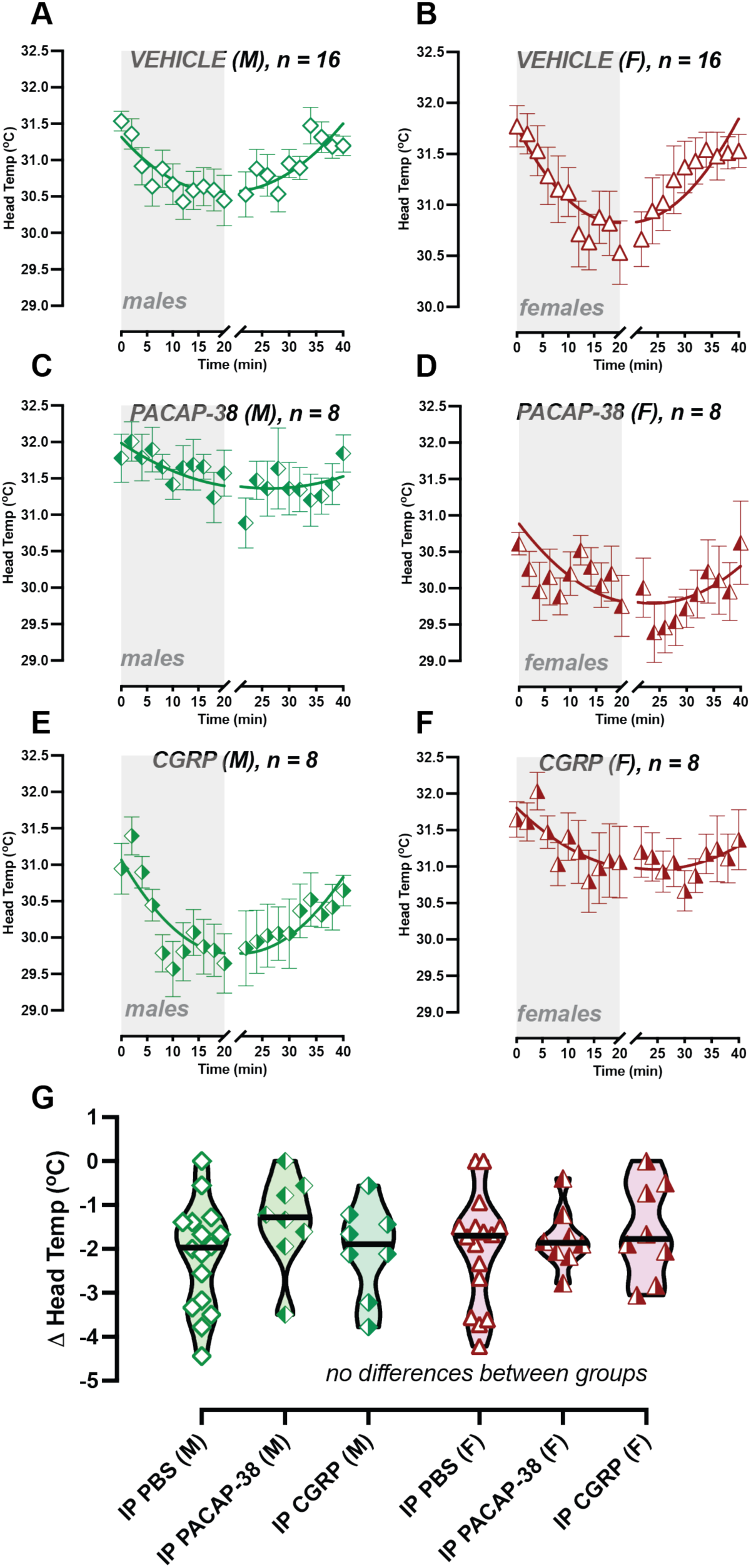
IP PACAP-38 and CGRP do not impact hypothermia magnitude. Head temperature profiles during testing are depicted for IP vehicle (**A and B),** IP CGRP (**C and D**), and IP PACAP-38 (**E and F**), with male graphs placed on left and females on right. Symbols are illustrated in the following scheme: green diamonds: male, red triangles: female, open symbol: IP vehicle, right-shaded symbol: CGRP, left-shaded symbol: PACAP-38. Sample sizes are depicted in legend. **G**. Delta head temperatures showed no differences in hypothermia between groups (Tukey post hoc). Hypothermia recovery is suggested to be prolonged after IP CGRP and IP PACAP-38 in females and males (data not shown, see **S1 Table 1**).

Mice exhibit hypothermia to provocative motion, and so we computed their delta head temperature, which represents the magnitude of their hypothermia, and compared the effects of IP vehicle, IP CGRP, and IP PACAP-38. We computed a 2-way mixed effects model examining the factors of Sex and Treatment but did not see a significant effect of either factor. Tukey multiple comparison’s test also showed no differences in delta head drops across treatment for males and females. We then examined the recovery from hypothermia using an X-intercept Quadratic Model after normalizing head temperatures (**S1 Table 1**). Mice treated with IP vehicle recovered from hypothermia at 19.2 minutes for females (R^2^ = 0.77) and 20.4 minutes for males (R^2^ = 0.70), which are comparable recovery times to previously published data [20], where we expect hypothermia recovery to resolve around 20 minutes after the end of provocative rotation. In mice treated with IP PACAP, hypothermia recovery was computed to be 26.7 mins for females (R^2^ = 0.34) and 28.0 for males (R^2^ = 0.47), and for IP CGRP, 27.0 mins for females (R^2^ = 0.61) and 21.00 mins for males (R^2^ = 0.70). We examined if the x-intercepts differ between data sets, and a significant difference was observed between x-intercepts for female Vehicle vs female PACAP-38 ( F(DF_n_, DF_d_) = 226.4 (1, 36), p < 0.0001), and female vehicle vs female CGRP (F(DF_n_, DF_d_) = 312.3 (1, 36), p < 0.0001). Additionally, a significant difference was observed between x-intercepts for male Vehicle vs male PACAP-38 (F(DF_n_, DF_d_) = 240.9 (1, 36), p < 0.0001), and male vehicle vs male CGRP (F(DF_n_, DF_d_) = 371.9 (1, 36), p < 0.0001). These findings suggest a significant disruption in tail vasodilation and a likely disruption in hypothermia recovery when mice are treated with IP PACAP-38, suggesting impairments to thermoregulatory control to provocative motion much like we have observed with IP CGRP, as shown in **Fig. 2G & H**.

### IP PACAP-38 increases postural sway in female C57BL/6J mice

To examine postural sway, mice were first tested after IP vehicle (n = 22F/18M) and then were divided into two groups, where one group was evaluated after IP CGRP (n = 14F/10M) and the other group for IP PACAP-38 (n = 8F/8M). We quantified their postural sway before and after the vestibular challenge (VC) by determining their center of pressure (CoP) represented as a 95% confidence ellipse area. We additionally computed the ellipse area’s major and minor axes. Three-way repeated measures ANOVAs (RM-ANOVA) analyzed CoP, major axes, and minor axes across the following three factors: Sex (male vs female), VC effect (pre and post VC), and Treatment (IP vehicle vs IP CGRP vs IP PACAP-38). Tukey multiple comparison’s test was used to further examine changes between groups.

Analysis of CoP showed effects of Sex (F (1, 74) = 67.06, p < 0.0001), a trending effect in Treatment (F (2, 74) = 2.725, p = 0.07), and no VC effect (F (1, 74) = 5.795e-006, p = 0.99), as shown in **Fig. 4A**. When we evaluated the interactions of these factors, we observed a significant interaction of Sex x Treatment (F (2, 74) = 7.326, p =0.001), and a significant interaction of Sex x Treatment x VC (F (2, 74) = 5.732, p = 0.005). In IP-vehicle treated females, we observed a change in the average CoP before and after the VC (adj. p = 0.008). In pre-VC measurements, females treated with IP CGRP exhibited larger CoP compared to their IP vehicle output (adj. p = 0.01), which resembles previous findings from our group [21]. Interestingly in pre-VC tests, females treated with IP PACAP-38 also exhibited larger CoP compared to their IP vehicle output (adj. p < 0.0001). Unlike females, males did not exhibit changes in their pre-VC CoP after IP CGRP or IP PACAP-38 (see **Fig. 4B**).

**Figure 4:**
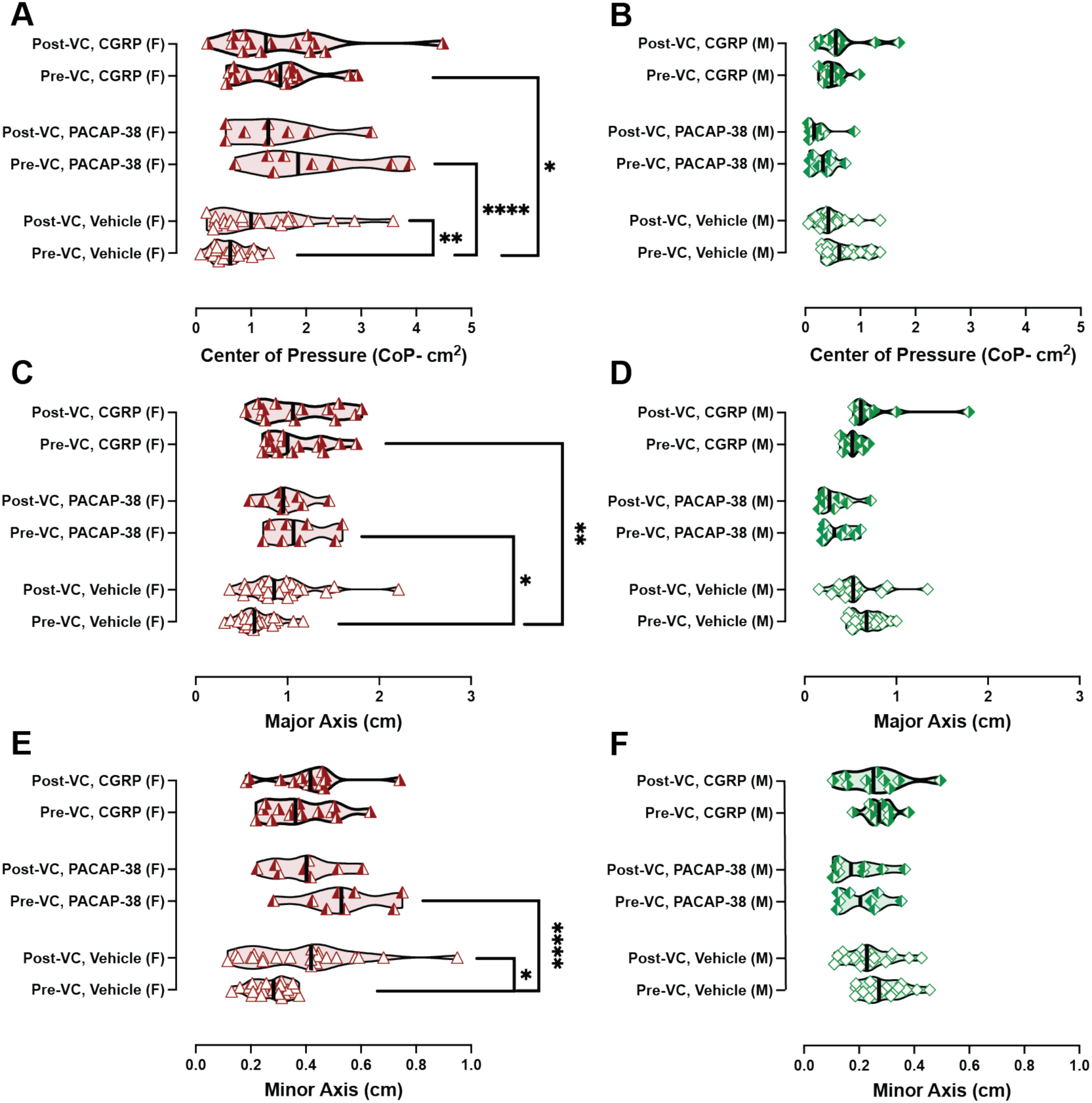
IP PACAP-38 and CGRP increase postural sway in female mice. Postural sway assessment involves assessing center of pressure 95% ellipse areas (**A and B**), major axes (**C and D**), and minor axes (**E and F).** Symbols are illustrated in the following scheme: green diamonds: male, red triangles: female, open symbol: IP vehicle, right-shaded symbol: CGRP, left-shaded symbol: PACAP-38. Sample sizes are depicted in legend. The vestibular challenge (5-minute orbital rotation, 125 rpm) led to (**A and E**) increased CoP and minor axes in IP-vehicle treated females. (**A and C)** In pre-VC testing, IP CGRP and IP PACAP-38 increased female CoP and major axes. Males did not seem affected by the neuropeptide treatments or vestibular challenge. Significance levels are indicated as follows: p < 0.05*, p < 0.01**, p < 0.001***

Analysis of major axes showed a sex effect (F (1, 74) = 70.23, p < 0.0001), a treatment effect (F (2, 74) = 4.804, p = 0.01), and no VC effect (F (1, 74) = 0.8482, p = 0.36). When we evaluated the interactions of these factors, we observed a significant interaction of Sex x Treatment (F (1, 74) = 70.23, p < 0.0001), and a significant interaction of Sex x Treatment x VC (F (2, 74) = 5.353, p = 0.007). In pre-VC measurements, females treated with IP CGRP exhibited longer major axes compared to their IP vehicle output (adj. p = 0.002), as shown in **Fig. 4C**. Females treated with IP PACAP-38 also exhibited longer major axes compared to their IP vehicle test (adj. p < 0.02).

In comparison to major axes, analysis of minor axes showed a Sex effect ((1, 74) = 47.98, p < 0.0001), no Treatment effect (F (2, 74) = 1.487, p = 0.23), and no VC effect (F (1, 74) = 0.45, p = 0.50), as shown in **Fig. 4E**. We additionally observed a significant interaction of Sex x Treatment (F (2, 74) = 6.636, p = 0.002), and of Sex x Treatment x VC (F (2, 74) = 5.19, p = 0.0078). In IP-vehicle treated females, we observed an increase in minor axes before and after the VC (adj. p = 0.03) as shown in **Fig. 4E**. In pre-VC minor axes for females, we did not observe an IP CGRP effect, but IP PACAP-38 had an effect (adj. p < 0.0001). Males did not show meaningful changes in their major axes or minor axes due to treatment or VC, as shown in **Fig. 4D & 4E.**

### IP PACAP-38 and a brief off-vertical axis rotation do not impact rotarod performance

Previously, we observed that IP CGRP led to deficits in rotarod performance when C57BL/6J mice are trained and tested on a mouse dowel (dowel radius = 1.5 cm) [22]. Thus, in this study, we assessed if these mice would exhibit rotarod deficits after IP PACAP-38, and if a brief but strong vestibular challenge (30 seconds of off-vertical axis rotation) would aggravate deficits. We did not observe rotarod deficits in mice tested after IP PACAP-38 and did not observe further changes after the OVAR in males and females (**Fig. 5B**).

**Figure 5:**
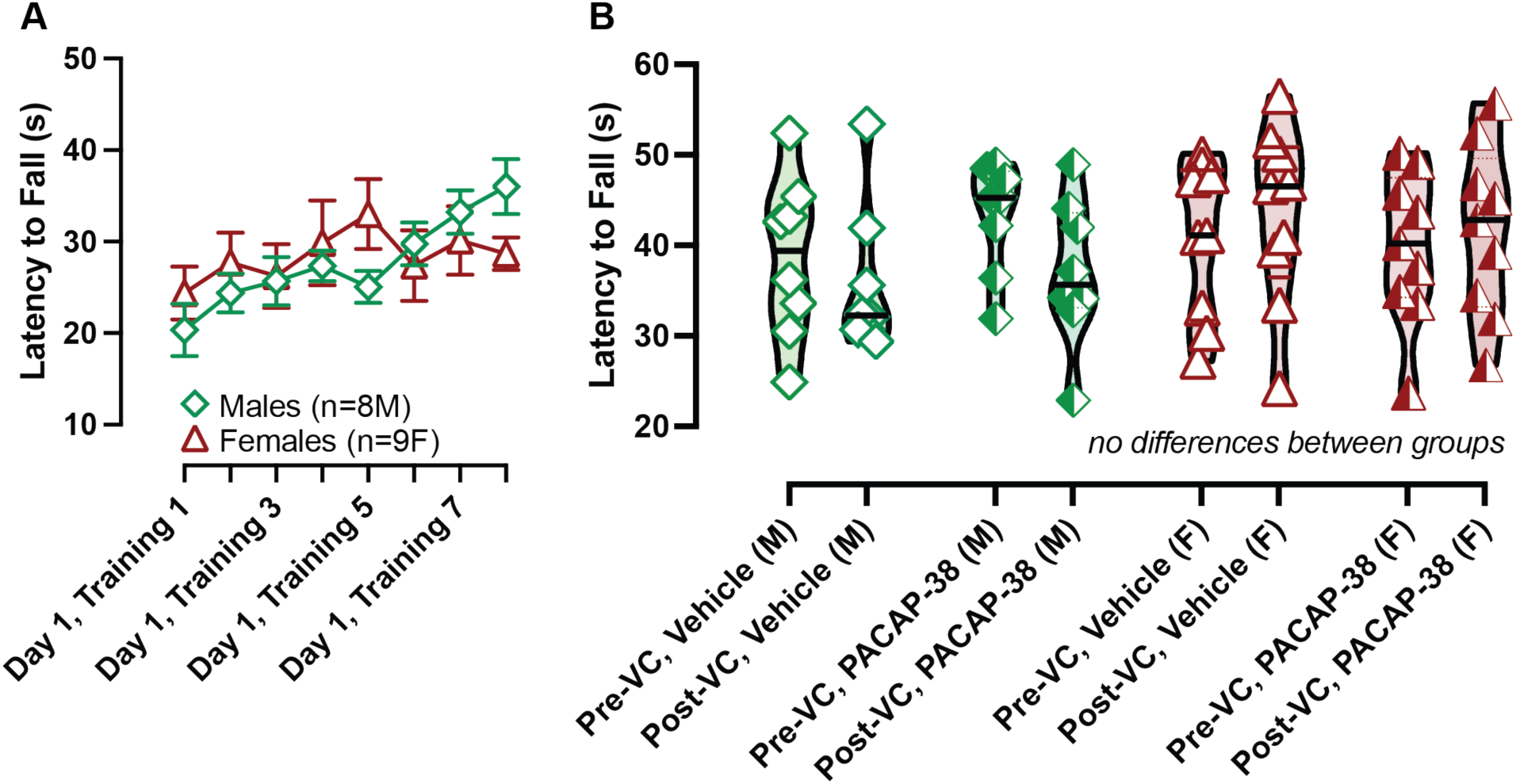
IP PACAP-38 does not impact rotarod using a mouse dowel. Mice were trained on day 1 with 8 trials (**A**) and tested for rotarod ability after IP vehicle (day 2) and IP PACAP-38 (day 3) (**B**). Max latency to fall (LTF) is plotted. Symbols are illustrated in the following scheme: green diamonds: male, red triangles: female, left-shaded symbol: PACAP-38. Sample sizes are depicted in legend. **A.** No differences were observed across sex, vestibular challenge, and PACAP-38 treatment.

### IP PACAP-38 significantly impacts balance beam performance

Mice were repeatedly assessed on balance beam assay after IP vehicle, PACAP-38, and CGRP injections (n = 5 F/ 5M), with each experiment capturing 3 trials per treatment and independently scored by two individuals. A two-way RM-ANOVA was used to assess the factors of biological sex (male vs female) and treatment (vehicle vs PACAP-38 vs CGRP) on time to traverse the balance beam in seconds (s), with uncorrected Fisher’s LSD test used for post hoc evaluation. Analysis of balance beam activity indicated effects of Treatment (F (2, 16) = 6.787, p = 0.007), but no effect of sex (F (1, 8) = 0.02, p = 0.9) or the interaction of these factors (F (2, 16) = 0.06, p = 0.94). Mice tested after IP vehicle administration clear the balance beam test quickly, as the times to traverse the balance beam, computed as a group mean + SEM, were 5.79 + 0.25 s for females and 7.292 + 0.34 s for males (**Fig. 6A**). In the vehicle control data, we observed tight confidence intervals and coefficient of variations (lower 95% CI, higher 95% CI; COV) for females (5.27 s, 6.3 s; 23.9%), and for males (6.61 s, 7.98 s; 25.2%), suggesting consistent and relatively stable performance on balance beam after IP vehicle. When mice were tested after IP PACAP-38, we observed significantly longer times to cross the balance beam (42.40 + 9.5 seconds for females, 43.45 + 8.7 seconds for males), and these times were longer than their vehicle control test for female (p = 0.018) and for male (p = 0.024). We also observed much wider confidence intervals and larger COVs for females (22.8 s, 62.0 s; 119%) and males (25.7 s, 61.2 s; 109.4%), suggesting a widespread in the range of observable outcomes (**Fig. 6B**). PACAP-38 injection significantly prolongs the time to traverse the balance beam in a significant proportion of mice, suggesting a balance or anxiety effect impacting performance in these mice.

**Figure 6:**
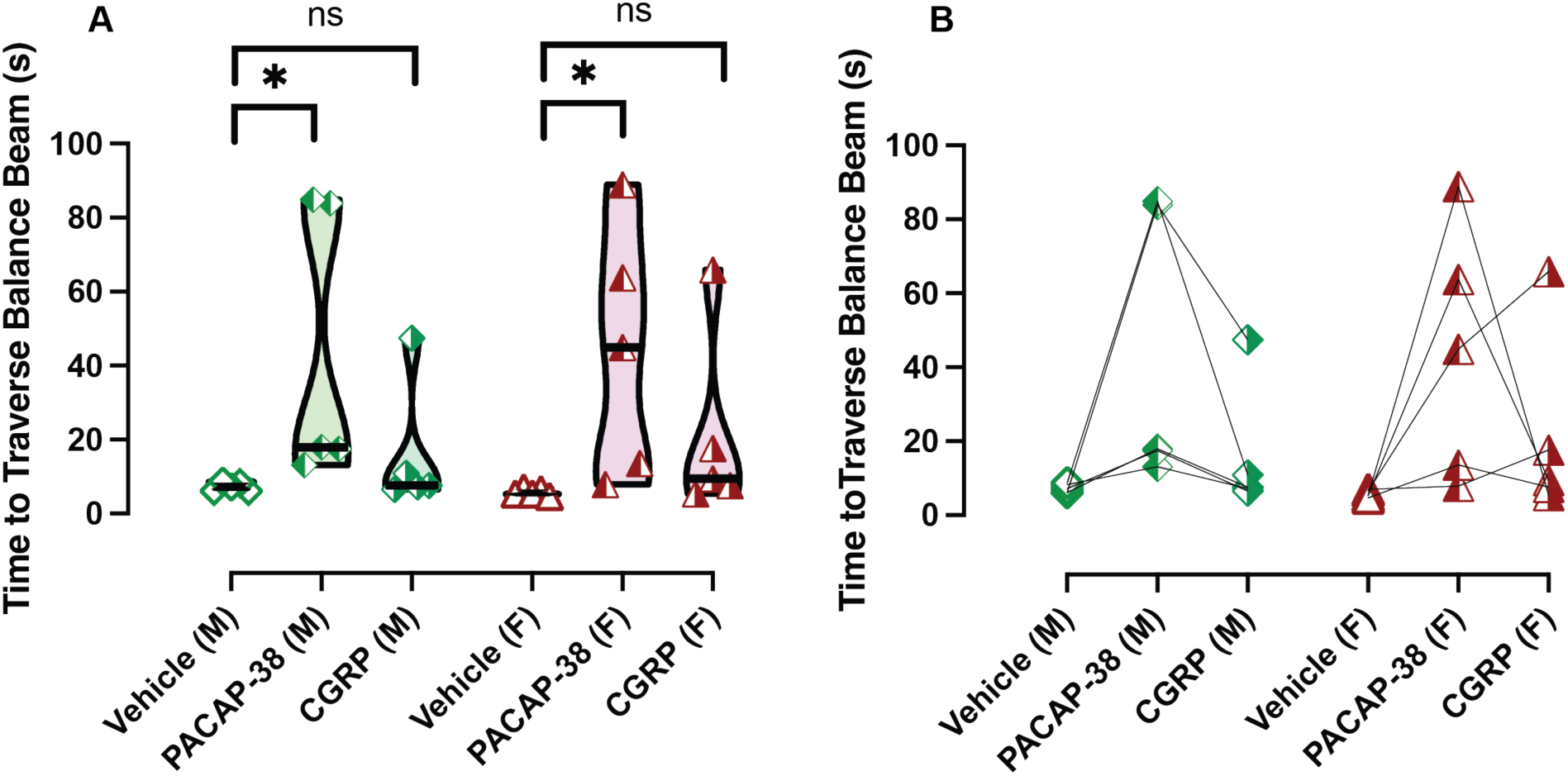
IP PACAP-38 disrupts balance beam behavior. Each symbol is the average of three trials per animal, verified by two independent scorers, for a given experiment (IP vehicle/CGRP/PACAP-38). Symbols are illustrated in the following scheme: green diamonds: male, red triangles: female, open symbol: IP vehicle, right-shaded symbol: CGRP, left-shaded symbol: PACAP-38. IP PACAP-38 significantly prolongs time to traverse balance beam in both sexes, while CGRP’s effects were not significant (**A**). Individual animal’s times are shown in **B**. Significance is listed as p < 0.05*.

Additionally, these mice were tested after IP CGRP. In a small number of males and females, we observed longer times to cross the balance beam, and computed the group mean + SEMs for mice to be 21.18 + 5.77 s for females and 15.94 + 5.2 s for males (**Fig. 6A**). Much like PACAP-38, we observed high confidence intervals and COVs in females (9.38 s, 33.0 s; 149.2%) and males (5.3, 26.6; 178.7%). The high variability also seen after IP CGRP testing suggests some mice are sensitized to CGRP’s effects and thus are impacted on the balance beam assay, but the majority of mice tested after IP CGRP did not show significant differences from their IP vehicle test, thus rendering IP vehicle vs IP CGRP group statistics to be not significant. Across treatments, we observed no significant difference between male and female performance. Both sexes show similar patterns of difficulty after injections of either neuropeptide. Since mice were repeatedly tested, a before/after plot is provided to show intra-animal changes (**Fig. 6B**).

## Discussion

In this study, we show that PACAP-38 disrupts motion-induced thermoregulation in mice, as IP PACAP-38 blunted tail temperature increases to provocative motion responses in males and females. IP PACAP-38 also increases postural sway in female C57BL/6J. Compared to their IP vehicle response, females displayed increased center of pressure (CoP) 95% ellipse areas and major axis lengths following IP PACAP-38, with notable differences observed before the vestibular challenge (VC). Importantly, these effects on postural sway were not observed in males. It is interesting to note that while females are significantly affected by systemic CGRP and PACAP-38, male mice are not affected. This parallels the finding that migraine and VM are significantly more prevalent in women than men, with some studies reporting a five-to-one female to male ratio [23]. IP PACAP-38’s effects on motion-nausea and sway mirror IP CGRP’s effects in this study and in our previous findings [20, 21]. However, unlike IP CGRP [22], we did not observe IP PACAP-38 to impact rotarod performance when mice were trained on a mouse dowel (r = 1.5 cm). On balance beam, IP PACAP-38 significantly prolonged the time required to traverse the balance beam in both male and female mice, indicating a potential impact on balance and/or anxiety. This increase in traversal time was accompanied by wider confidence intervals and high coefficients of variation in both sexes, suggesting a wide range and differential sensitivity even within the C57BL/6J strain. In contrast, IP CGRP did not significantly affect balance beam activity, but we did observe a few mice to experience longer traversal times compared to their vehicle test.

Our study demonstrates that intraperitoneal (IP) administration of PACAP-38 disrupts mouse surrogate behaviors for motion-induced nausea. The surrogate measure of motion-induced nausea is the transient tail temperature spike upon rotation. It is puzzling that both systemic CGRP and PACAP cause this transient spike in tail temperature to decrease or be eliminated rather than to increase if purely a vasodilatory function. We believe this tail temperature spike to vehicle is the animal using its tail to actively resist rotation, whereas after IP CGRP or PACAP-38 the animal no longer mounts a response to rotation and is passively enduring the rotation.

We also found that IP PACAP-38 disrupts other mouse surrogate behaviors for balance. PACAP can cross the blood-brain barrier as well as act peripherally in brainstem and inner ear [24]. We don’t know where PACAP-38 is acting in the CNS though could be a combination of sites as PACAP and CGRP are involved in modulating neuronal activity within the CNS and efferent pathways of the vestibular system, including pathways related to sensory processing and motor responses [25].

PACAP-38 activates a G-protein-coupled receptor (GPCR) family consisting of the PAC1 receptor and the vasoactive intestinal peptide receptors (VPAC1/VPAC2). PACAP peptides (PACAP-38 and -27) preferentially activate the PAC1 receptor, but VPAC1 and VPAC2 receptors can be activated by both PACAP and vasoactive intestinal peptide (VIP) [26]. A previous study showed that IP PACAP-38 induces light aversion in the CD1 mice, and using RNA-seq and gene expression analyses, observed that in mice sensitive to PACAP-38’s effects (responder mice), IP PACAP-38 led to increased pituitary glycoprotein hormones and receptor RNAs in the trigeminal ganglia of male mice and elevation of *Trpc5* ion channel RNA in male and female mice [11]. The elevation of these RNAs is suggested to impact nociception, chemosensory avoidance [27], and peripheral sensitization [28].

To date, there is limited research on PACAP’s role in vestibular sensory processing and related complications like motion sickness. PACAP-sensitive fibers are found in the dorsal and ventral horn of the spinal cord, and PACAP-containing cell bodies are found in the brainstem, hypothalamus, and area postrema [29]. The area postrema – also known as the chemoreceptor trigger zone (CTZ) – regulates nausea sensation and lies outside of the blood brain barrier, able to detect toxins in blood and in the CSF [30]. The CTZ contains PACAP-positive fibers and PACAP-binding sites [29, 31, 32]. Moreover, a subset of brainstem neurons in the nuclei of the solitary tract and area postrema control multiple aspects of sickness behaviors [33], yet others found that surrogates of nausea and vomiting to general anesthetics involves area postrema in rats [34]. However, other studies found that lesions to area postrema did not affect motion-sickness in rats or cats, so perhaps lesions to area postrema eliminate nausea to toxins and not to motion [35, 36].

Both PACAP and CGRP and their receptors are found in the trigeminal system [37, 38]. In fact in mice that have chemical activation of trigeminal system y nitroglycerin only PACAP (+/+) mice showed increases in c-Fos expressing cells and showed changed in photophobia light-aversion behaviors that were absent in PACAP (-/-) null mice [39]. Both CGRP and PACAP are also present in the efferent vestibular system (EVS) of the brainstem. This EVS system can modify vestibular signal at the peripheral vestibular organs. Therefore, this efferent feedback system maybe involved in development of motion-sickness symptoms that are enhanced by systemic CGRP or PACAP [25]. In addition, we have shown CGRP-positive fibers in the crista and otolithic vestibular organs and others have shown PACAPS present in 2^nd^ and 3^rd^ order sensory neurons of the cochlear and vestibular systems [40–42].

CGRP blocking drugs appear to be ineffective in blocking PACAP-38’s effects in rodent models for light aversion, suggesting these peptides operate independently, and thus its of future interest to evaluate PACAP and CGRP blockade using the behavioral assays in this study [43]. Interestingly, in migraine sufferers that responded to anti-DGRP monoclonal antibodies, after treatment they show reduced functional connectivity between primary somatosensory and motor cortices, insular, and other occipital and temporal-occipital areas; whereas the non-responders showed no connectivity differences from before treatment [44]. Recent studies and clinical trials have found anti-CGRP based migraine medications to be effective treatments for VM in 65-70% of cases [18, 19].

The PACAP antagonist, LU AG09222, is currently being assessed in a Phase 2 clinical trial for migraine relief (clinical trial identifier: NCT06323928), and is currently ongoing, but there are no studies evaluating PACAP antagonism for VM therapy. Our preclinical findings warrant an opportunity to evaluate PACAP blockade as a therapy for VM patients, as many VM patients are non-responders to both triptans and anti-CGRP medications [18].

Overall, our findings highlight the influence of PACAP-38 on motion-induced nausea, balance beam imbalance, and postural sway, which may have implications for understanding migraine and vestibular migraine pathophysiology and developing targeted treatments.

## Acknowledgements

We would like to thank Najla Silmi for contributions to analysis of balance beam data.

## Figure and Table Legends

**S1 Table 1.** To evaluate recovery from hypothermia, an x-intercept quadratic model was used, with a constraint imposed to detect x-intercepts after time t = 20 minutes (when provocative motion is turned off). Recovery period is calculated by taking the x-intercept (minutes) and subtracting 20 minutes (duration of provocative rotation from t = 0 to t = 20 mins).

| Analyses from X-Intercept Quadratic Model across sex and treatment for Head Temperature Profiles |  |  |  |  |  |  |
| --- | --- | --- | --- | --- | --- | --- |
| Variables | Vehicle (F) | Vehicle (M) | PACAP-38 (F) | PACAP-38 (M) | CGRP (F) | CGRP (M) |
| **Best-fit values** |  |  |  |  |  |  |
| *slope1* | 0.096 | 0.090 | 0.039 | 0.065 | 0.062 | 0.116 |
| *slope2* | 0.002 | 0.002 | 0.0008 | 0.001 | 0.001 | 0.003 |
| *X-intercept (mins)* | 39.21 | 40.36 | 47.97 | 46.67 | 46.95 | 41 |
| *Recovery period (mins)* | 19.21 | 20.36 | 27.97 | 26.67 | 26.95 | 21 |
| **95% CI (profile likelihood)** |  |  |  |  |  |  |
| *slope1* | 0.08 to 0.11 | 0.07 to 0.11 | 0.02 to 0.06 | 0.029 to 0.09 | 0.04 to 0.08 | 0.09 to 0.14 |
| *slope2* | 0.002 to 0.003 | 0.001 to 0.003 | 0.0002 to 0.002 | 0.0003 to 0.002 | 0.0007 to 0.002 | 0.002 to 0.004 |
| *X-intercept* | 37.22 to 42.02 | 37.95 to 44.08 | 39.80 to 110.2 | 40.06 to 78.44 | 41.46 to 61.39 | 38.33 to 45.29 |
| **Goodness of Fit** |  |  |  |  |  |  |
| *Degrees of Freedom* | 18 | 18 | 18 | 18 | 18 | 18 |
| *R squared* | 0.768 | 0.697 | 0.470 | 0.344 | 0.615 | 0.702 |
| *Sum of Squares* | 0.65 | 0.72 | 0.78 | 1.54 | 0.76 | 1.33 |
| *Sy.x* | 0.19 | 0.2 | 0.2 | 0.29 | 0.21 | 0.27 |

